# Heat stress response and transposon control in plant shoot stem cells

**DOI:** 10.1101/2023.02.24.529891

**Authors:** Vu Hoang Nguyen, Ortrun Mittelsten Scheid, Ruben Gutzat

**Affiliations:** Gregor Mendel Institute of Molecular Plant Biology, Austrian Academy of Sciences, Vienna Biocenter (VBC), 1030 Vienna, Austria

**Keywords:** Heat stress, DNA methylation, RdDM, Shoot apical meristem, Stem cells, Transposon

## Abstract

Post-embryonic plant development must be coordinated in response to and with environmental feedback. Development of above-ground organs is orchestrated from stem cells in the center of the shoot apical meristem (SAM). Heat can pose significant stress to plants and induces a rapid heat response, developmental alterations, chromatin decondensation, and activation of transposable elements (TEs). However, most plant heat-stress studies are conducted with whole plants, not resolving cell-type-specific responses. Heat stress consequences in stem cells are of particular significance, as they can potentially influence the next generation. Here we use fluorescent-activated nuclear sorting to isolate and characterize stem cells after heat exposure and after a recovery period in wild type and mutants defective in TE defense and chromatin compaction. Our results indicate that stem cells can suppress the heat response pathways that dominate surrounding somatic cells and maintain their developmental program. Furthermore, mutants defective in DNA methylation recover less efficiently from heat stress and persistently activate heat response factors and heat-inducible TEs. Heat stress also induces epimutations at the level of DNA methylation, and we find hundreds of DNA methylation changes three weeks after stress. Our results underline the importance of disentangling cell type-specific environmental responses for understanding plant development.

## 1. Introduction

Transposable elements (TEs) are mobile genetic elements that can replicate independently of the host genome. Transposition of TEs within the host genome can result in mutations or changes in gene expression (Dubin et al., 2018; Lisch, 2013). Since the seminal discovery of TEs in maize by Barbara McClintock (McClintock, 1956), transposons have been investigated intensively, particularly after the advancement of sequencing technology (Cain et al., 2020). Studies with somatic tissues (Guo et al., 2021; Sun et al., 2020) and gametophytes (Slotkin et al., 2009; Olmedo-Monfil et al., 2010) showed that transposon control could differ between somatic and germline cells. However, as the germline separation in plants is not so clear and debated (Lanfear, 2018), these differences need to be investigated in more detail. Recent evidence shows that populations of plant stem cells of the shoot apical meristem (SAM) are similar in some respect to germline cells (Bradamante et al., 2022; Nguyen and Gutzat, 2022) and are hubs for TE activity. As SAM stem cells give rise to all post-embryonic above-ground organs, including the reproductive cells, these cells are “of interest” for TEs to increase their copy number vertically over generations.

TEs are classified into two groups. Class I or retrotransposons amplify through a “copy-and-paste” mechanism, using RNA replication intermediates and reverse transcriptase (Lisch, 2013). After reverse transcription, double-stranded DNA is produced to form linear or circular extrachromosomal DNA (Fan et al., 2022). The extrachromosomal DNA is inserted into the genome by integrases. Class I includes long terminal repeat retroelements (LTR), non-LTR retroelements, and short interspersed nuclear elements (SINEs). Class II TEs or DNA transposons employ a “cut-and-paste” mechanism to transpose (Lisch, 2013) via a transposase encoded in their genome responsible for excision and insertion. Class II TEs encompass miniature inverted-repeat transposable elements (MITEs), including degenerate non-autonomous copies that nevertheless can be mobilized in the presence of autonomous copies (Feng, 2003). Helitrons are also classified as DNA transposons, although their transposition via a rolling circle mechanism is slightly different (Kapitonov and Jurka, 2001; Thomas and Pritham, 2015).

TEs occupy a large fraction of plant genomes (Charles et al., 2008; Huang et al., 2012; Jiao et al., 2017) and can amplify when the epigenetic control is impaired (Ito et al., 2011; Marí-Ordóñez et al., 2013). Epigenetic TE silencing mechanisms include DNA methylation, heterochromatic histone modifications, and higher-order compaction of chromatin (Liu et al., 2022). A crucial regulator of DNA methylation is the chromatin remodeler DECREASE IN METHYLATION 1 (DDM1), discovered decades ago by the loss of DNA methylation at satellite repeats in the mutant (Vongs et al, 1993). Loss of DDM1 allows transcription and reinsertion of different transposons (Higo et al., 2012; Hirochika et al., 2000; Kato et al., 2004). DDM1 exerts its role in DNA methylation likely by facilitating chromatin access to DNA methyltransferases such as METHYLTRANSFERASE 1 (MET1) and CHROMOMETHYLASE 2/3 (CMT2/3). Loss of DDM1 results in an almost complete reduction of DNA methylation in all sequence contexts (Zemach et al., 2013). DNA methylation at CHG and CHH sites of TEs is installed by RNA-directed DNA methylation (RdDM). In this pathway, a plant-specific polymerase POLYMERASE IV (POLIV) with an affinity for methylated DNA produces short ∼40 nt-long transcripts (Zhai et al., 2015). These transcripts are processed by RNA-dependent RNA polymerase 2 (RDR2) and DICER 3 (DCL3) into 24 nt-long siRNAs and loaded onto nuclear ARGONAUTE (AGO) proteins. Together with the AGOs, the siRNAs guide the RNA-induced silencing complex (RISC), which includes methyltransferases DOMAIN REARRANGED METHYLTRANSFERASE 1/2 (DRM1/2), to complementary transcripts of another plant-specific polymerase, POLYMERASE V (POLV). DRM1/2 methylates DNA asymmetrically in a CHH context (H stands for any base but G) The analysis of various mutants indicates that this RNA-based pathway of TE control (Wendte and Pikaard, 2017) determines mainly methylation of short TEs on chromosome arms, whereas DDM1 has a more prominent role for DNA methylation on long pericentromeric TEs (Zemach et al., 2013).

While the epigenetic control of TEs is usually tight and strongly determined by the components described above, biotic and abiotic stress can transiently suspend TE silencing and trigger their activity (Dubin et al., 2018). One example is the Arabidopsis retrotransposon ONSEN which becomes transcribed at high temperatures (Tittel-Elmer et al., 2010) and forms extrachromosomal DNA (Cavrak et al., 2014). No new insertions were found in wild-type plants and within the experimental time frame, but ONSEN proliferated in *poliv* and other RdDM mutants after heat stress (Ito et al., 2011; Hayashi et al., 2020). Analysis of the insertion pattern along the inflorescence revealed that the insertion patterns differed between seeds obtained from individual flowers but were similar within the progeny from the same flower (Ito et al., 2011). Therefore, transposition happened before gamete specification in somatic tissue, giving rise to the germline. Heat stress also has many other effects on plant development, like accelerated flowering (Balasubramanian et al., 2006), which must be controlled by the development of the SAM and the stem cells therein. Furthermore, there is evidence that plants have evolved protection mechanisms against heat stress involving factors in the stem cells (Olas et al., 2021). Therefore, in this study, we aimed to understand the heat stress response of SAM stem cells compared to somatic cells. To consider the two different pathways contributing to the epigenetic control of TEs described above, we analyzed stem cells of wild-type plants and *ddm1* and *poliv* mutants for potential changes in DNA methylation triggered by heat stress and for their maintenance beyond the stress period.

We used the heat-dependent transcriptional activation of ONSEN to establish sublethal but effective heat stress conditions. We then performed fluorescence-activated nuclei sorting (FANS) (Gutzat and Mittelsten Scheid, 2020) to collect stem cells for transcriptome and methylome analysis from plants shortly after the stress and after a recovery period.

Our results uncover tissue- and genotype-specific heat stress responses. We find that stem cells can maintain developmental transcriptional programs that are down-regulated when analyzing whole seedlings. Furthermore, we show that both DNA methylation mutants are hypersensitive to heat treatment and that *POLIV* and *DDM1* are critical in suppressing the heat stress response, including heat response factors and heat-induced transposons during recovery. We also detect heat-induced DNA methylation changes in stem cells, especially at CHG sites, in accordance with their dynamic methylation pattern in these cells during development (Gutzat et al., 2020).

## 2. Materials and methods

### Plant material and growth conditions

*Arabidopsis thaliana* ecotype *Col-0* was used for all experiments. The stem cell reporter *pCLV3::H2BmCherry* is described in (Gutzat et al., 2020) and was crossed with *poliv* (*nrpda-3* - SALK_128428) and *ddm1* (*ddm1-10* - SALK_093009). Plants were grown either *in vitro* on GM medium or soil, with 16/8 h light/dark cycles. Control plants were consistently grown at 21°C; for heat exposure, we applied 37°C and 44°C for 24 h or 48 h.

### DNA extraction

To prepare DNA for methylation analysis by bisulfite-sequencing, 5000 FANSed nuclei were used to extract DNA with the Quick-DNA microprep kit (Zymo Research #D3020).

For Southern blots, DNA was extracted from 300 mg of above-ground tissue of seedlings with CTAB adapted from (OPS Diagnostics CTAB protocol for isolating DNA).

### RNA extraction

Ten dissected seedlings or ten inflorescence meristems were used for trizol RNA extraction adapted from (Rio et al., 2010).

### Southern blots for the detecting of ONSEN extrachromosomal DNA

For Southern blot detection, 10 µg of genomic DNA was restricted by *Hpa*II (ThermoFisher # ER0511) overnight. After enzyme inactivation (95°C for 5 min) DNA was precipitated with sodium acetate and ethanol and loaded on a 1% TAE agarose gel. The gels were rinsed with 250 mM HCl for 10 min, then with denaturation solution (500 mM NaOH, 1.5 M NaCl) for 30 min, and incubated in neutralization buffer (0.5 M Tris, 1.5 M NaCl, 1 mM EDTA) for 30 min.

DNA was transferred onto a Hybond NX membrane (Merck #GERPN203T) and cross-linked by a UV linker (Stratagene UV Stratalinker 2400).

The probe was generated by PCR amplification of the ONSEN sequence with forward primer TCTAGAATCATCTTCCACCTCCTTA and reverse primer ATCCTTGATAGATTAGACAGAGAGCT. The resulting amplicon was labelled with α-^32^P-dCTP using the labeling kit (Aligent #300385).). The membranes were hybridized with the denatured probe at 65°C overnight in hybridization buffer (0.4 M Na_2_HPO_4_, 0.1 M NaH_2_PO_4_, 20% SDS, 0.5 mM EDTA) and exposed to a phosphor screen for one day and imaged by phosphor-imager (Amersham Typhoon).

### Fluorescence-activated nuclei sorting (FANS)

FANS was described before (Gutzat and Mittelsten Scheid, 2020).

### mRNA and bisulfite library preparation and sequencing

From FANS-sorted material, bulks of 100 nuclei for each replicate were used for library preparation. Smart-seq v2 (seedling, IM, stem cell D28) and v3 (stem cell D7) library preparation and sequencing were performed by the Next Generation Sequencing Facility (Vienna BioCenter Core Facility) NovaSeq SP SR100. For bisulfite sequencing, libraries were prepared with the Pico Methyl-Seq prep kit (Zymo Research #D5456) and sequenced by the Next Generation Sequencing Facility (Vienna BioCenter Core Facility) NovaSeq SP SR100.

### RNA-seq analysis

Raw bam files were converted to fastq by bedtools (v2.27) (Quinlan and Hall, 2010). The fastq files were used as input for the nextflow rnaseq pipeline (v21.02.0) (Ewels et al., 2020) with additional parameter “--clip_r1 19 --clip_r2 9 --three_prime_clip_r1 5 --three_prime_clip_r2 5”. Output count files were used to analyze differential gene and TE expression by DESeq2 (Love et al., 2014) with filtering out of loci with less than 5 reads per genotype on avarage and with the cutoff of p.adj. < 0.05, log2 fold change > |1|. Gene ontology enrichment was performed using the PANTHER classification system (geneontology.org). Visualization of the data was done with R using the packages tidyverse, ggplot2 (Wickham, 2016), pheatmap (Kolde, 2018), and chromplot (Oróstica and Verdugo, 2016).

### Bisulfite-seq analysis

Raw bam files were converted to fastq by bedtools (v2.27) (Quinlan and Hall, 2010). Adaptor were removed with trim galore (v0.6.2) (Krueger et al., 2021) with parameter --clip_r1 13 -- three_prime_clip_r1 12. Genome alignment was performed with bismark (v0.22.2) (Krueger and Andrews, 2011) with bowtie2. Output files were deduplicated by deduplicated_bismark (bismark v0.22.2). Methylation level was obtained from methylpy call-methylation-state (v1.2.9) (Schultz et al., 2015). Differential methylated regions were obtained using a two-state Hidden Markov Model (HMM) method on nextflow methylscore (v21.10.06) (Hüther et al., 2022).

## 3. Results

### 3.1. Identification of heat stress conditions

To establish heat stress conditions that are non-lethal but would induce a maximum of TE activation and potentially DNA methylation changes, we tested different temperature regimes and culture conditions, quantifying ONSEN activity as an indicator of response. Heat-activated ONSEN forms extrachromosomal DNA from retrotranscribed RNA, which can be quantified by Southern blot analysis (Ito et al., 2011; Cavrak et al., 2014). Plants were cultured on soil or *in vitro* and exposed to varying temperatures, as described in Figure 1a,b. Plants grown *in vitro* displayed a much higher abundance of ONSEN extrachromosomal DNA than soil-grown plants (Figure 1b). This is likely due to increased humidity and reduced cooling by transpiration within the culture vessels. We also tried initial cold treatment, as this has been used in previous studies (Ito et al., 2011). In our growth conditions, cold treatment did neither induce extrachromosomal ONSEN nor enhance ONSEN response to subsequent heat treatment (Figure 1b). Maximum ONSEN abundance was observed after heat stress at 37°C for two days (Figure 1b). The non-transgenic Col-0, as well as the line with the SAM stem cell reporter *pCLV3::H2BmCherry* (Gutzat et al., 2020), hereafter named *wt*, survived this harsh treatment well, but unexpectedly, this stress regime was lethal for seedlings of the DNA methylation mutants *poliv* and *ddm1* (Figure 1c). Both mutants showed necrotic spots on leaves and cotyledons after one day at 37°C but survived in the recovery period (Figure 1d). To allow inclusion of the mutants in the experiments, we chose *in vitro* growth of the plants and exposure for one day at 37°C, providing strong ONSEN induction but low lethality of all genotypes.

**FIGURE 1.**
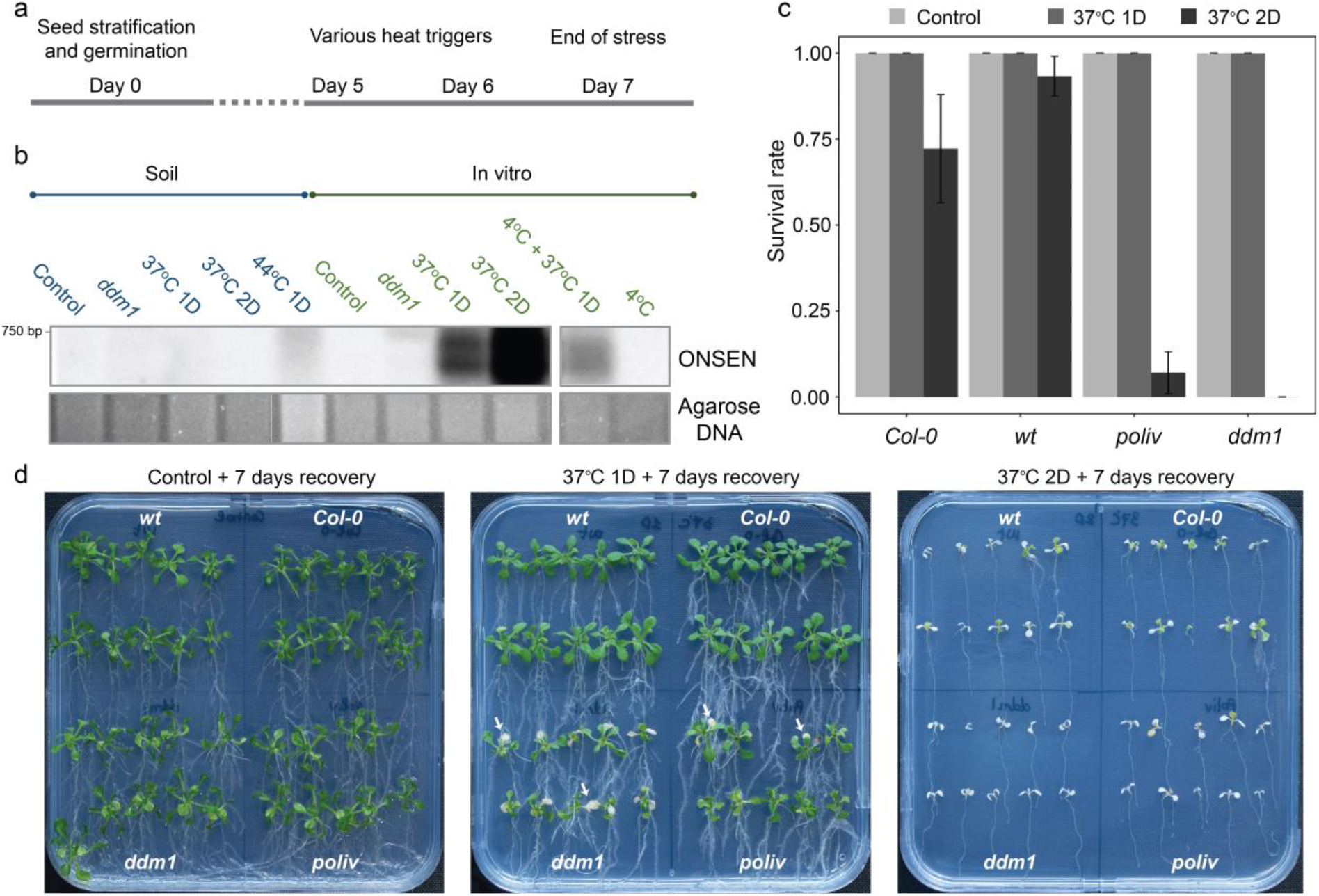
Establishing heat stress conditions. (a) Experimental setup for testing different heat stress conditions. (b) Southern blot probing for ONSEN extrachromosomal DNA. Culture medium, temperature, and exposure time for each lane are indicated. All samples are from wt plant except *ddm1*. The full blots are shown in Supplementary Figure 1. (c) Survival rates of *Col-0, wt* (*pCLV3::H2BmCherry*), *poliv*, and *ddm1* plants after seven days of recovery at 21°C, N = 3 (30 plants total/genotype). (d) Images of plants seven days after heat exposure.

### 3.2. Gene expression changes induced by heat

We aimed to understand (i) whether gene expression under heat stress differs between stem cells and somatic tissue; (ii) if and how these changes are influenced by DNA methylation; (iii) and whether changes in gene expression or DNA methylation induced by heat stress persist in stem cells, with the potential to become heritable.

To address these questions, we performed heat and mock treatment in *wt, poliv*, and *ddm1* plants, all containing the reporter labelling the nuclei of SAM stem cells. We collected above-ground tissue from seedlings and stem cell nuclei using FANS right after heat stress (D7). For *wt*, we also collected non-stem cell nuclei, representing nuclei of cells from above-ground tissue. The remaining seedlings were transferred to soil and grown for an additional 21 days when they started to flower (D28). At this time point, we again collected stem cell nuclei by FANS and hand-dissected apices with inflorescence meristems.

Collecting stem cell nuclei after heat stress was challenging, as they were more sensitive to the extraction procedure than mock-treated samples, and heat-stressed tissues displayed more autofluorescence. This resulted in a very low yield of intact nuclei in general and even less for stem cell nuclei. However, as we could obtain genome-wide representative transcripts from only 50 nuclei in combination with smart-seq3 single-cell sequencing (Bradamante et al., 2022), we used bulks of 100 nuclei as samples for FANSed material, in combination with smart-seq3 single-cell sequencing for all mRNA-seq samples.

Principal component analysis of the mRNA data showed a clear grouping of the samples (Figure 2). PC1 separated *ddm1* from *wt* and *poliv* samples, independently of treatment or tissue type. PC2 separated mostly tissue types and stress conditions. Noticeably, upon heat stress, seedlings (+Sd) became more similar to FANSed stem cell (+SC7) and non-stem cell (+nSC7) samples, but stem cell samples (+SC7) more different to non-stem cell samples (+nSC7). This suggests that a standard set of genes is changed by high temperature but also indicates specific responses in stem cells. Furthermore, at the recovery stage, heat- and mock-treated stem cell nuclei (+SC28; -SC28), as well as inflorescence meristems (+IM28; -IM28), overlapped, implying that most genes had returned to a similar expression state as before stress.

**FIGURE 2.**
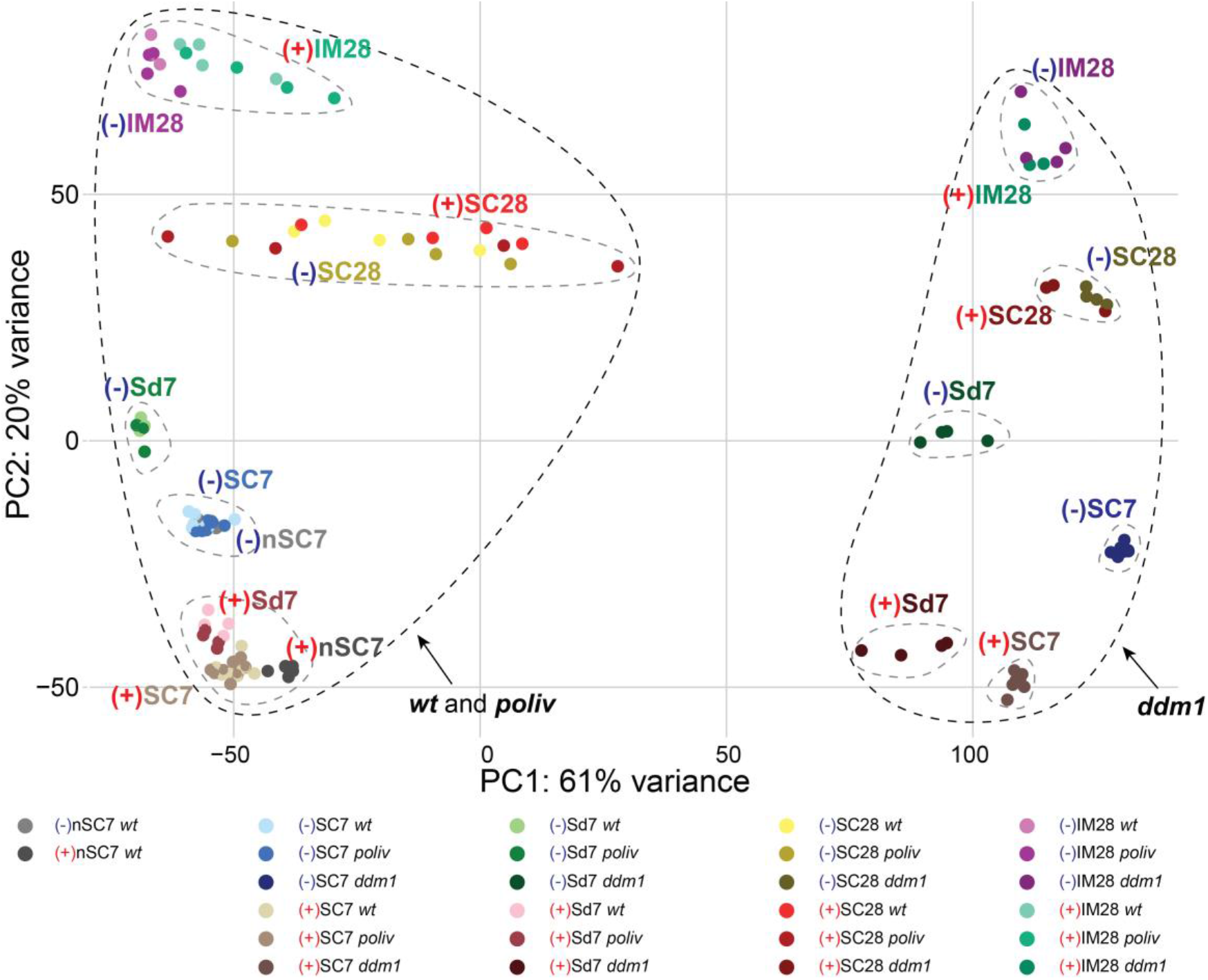
Principal component analysis shows clusters within the mRNA datasets. Arrows indicate two groups. Nomenclature: (-): control; (+): heat stress; nSC: non-stem cells; SC: stem cells; C7D: control at D7; H7D: heat stress at D7; Sd: seedlings; C28D: control plant at D28; H28D: recovered from heat stress at D28; IM28: inflorescence meristems.

For all genotypes, the gene expression differences between stem cells and the surrounding tissue (SC7 versus Sd7) were more prominent at the later developmental stage (SC28 versus IM28). Interestingly, a slight separation persisted 21 days after heat treatment for the inflorescence meristems (+IM28 versus -IM28), which could indicate some long-lasting transcriptome changes. However, we could not observe this in the *ddm1* samples, and all *ddm1* samples are separated from the other two genotypes, likely due to the general deregulation of gene expression in this mutant.

Taken together, the results show that the impact of severe heat stress on gene expression can reduce tissue-specific differences but cannot override the substantial *ddm1*-specific differences, connected with reduced DNA methylation in all sequence contexts and severe chromatin decompaction already before heat stress (Probst et al., 2003). Furthermore, the lack of asymmetric DNA methylation in heterochromatin, characteristic of *poliv*, does not influence a global transcriptional response. Nevertheless, comparing the differences between stem cells and somatic cells right after heat stress and between *wt, poliv*, and *ddm1* will be informative in understanding the mutants’ increased heat sensitivity and the dependence of DNA-specific responses on DNA methylation.

### 3.3. Heat response differences in stem vs. somatic cells

We investigated the gene expression differences in stem cell vs. non-stem cell nuclei directly after heat stress in more detail. Expression of 94 reference genes for stressed shoots (Czechowski et al., 2005) showed no significant differences (Figure 3a), excluding systematic differences between the samples. *AGO5* and *AGO9* are specifically expressed in stem cells (Bradamante et al., 2022), and the expression of *CLV3, AGO5*, and *AGO9* confirmed high enrichment in stem cells (Figure 3a). *AGO5* and *AGO9* transcripts were even elevated in stem cell nuclei after heat stress (Figure 3a). In addition, the number of stem cell-specific differentially expressed genes (DEGs: Wald test FDR < 0.05 and log2 fold change > |1|) was more than doubled by heat stress (Figure 3b), demonstrating a strong stem cell-specific heat response. After heat treatment, we found 515 genes up- and 423 genes down-regulated exclusively in stem cells (Figure 3c). Analyzing the expression pattern of these stem cell-specific DEGs revealed five clusters with a strong difference between stem- and non-stem cells (Figure 3d).

**FIGURE 3.**
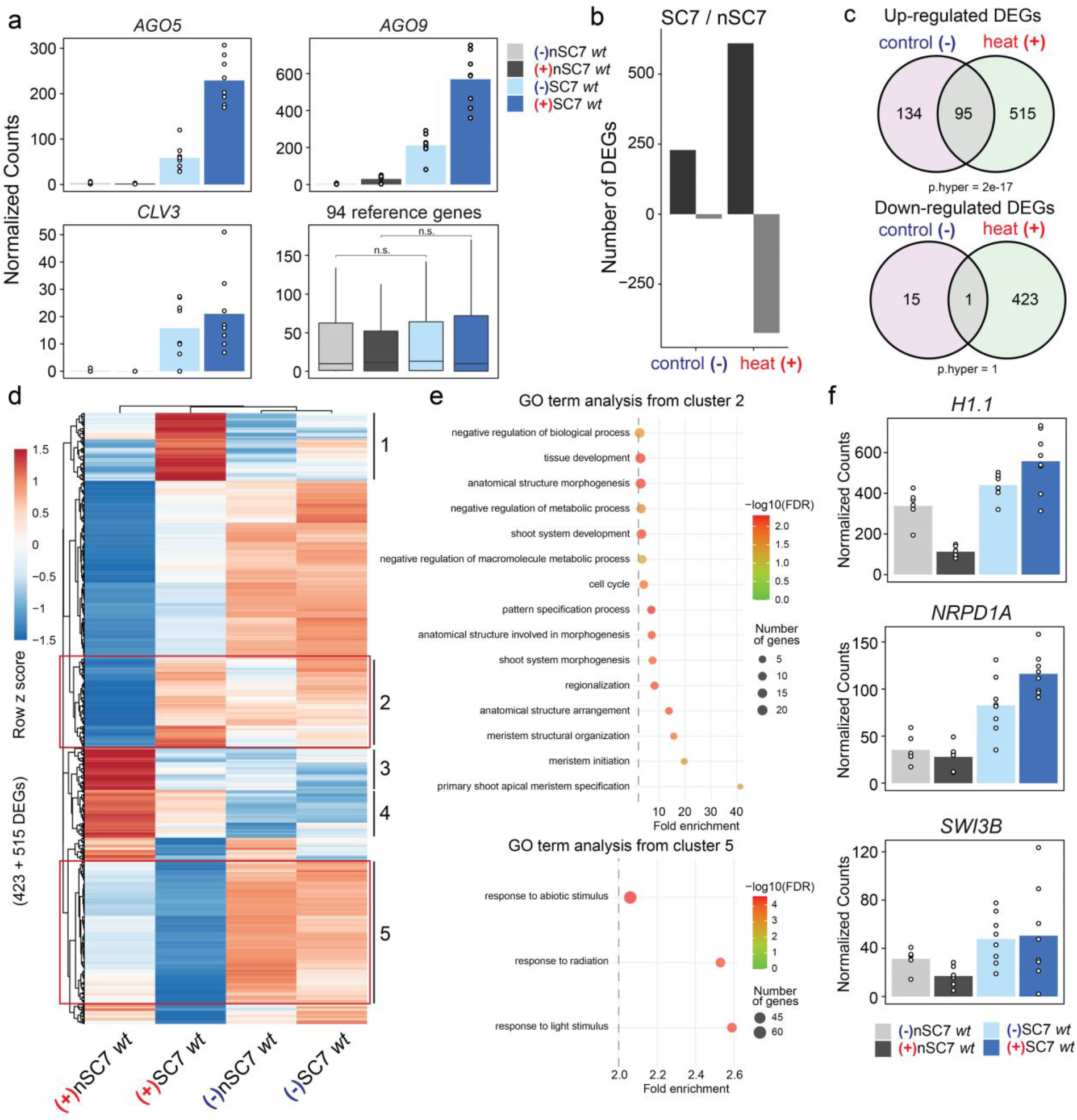
Gene expression analysis confirms stem cell-specific gene expression even after heat stress. (a) Normalized counts of reads for stem cell markers: *AGO5, AGO9, CLV3*, and the average of 94 reference genes. (b) Number of differentially expressed genes (DEGs) by pairwise comparison SC7 vs. nSC7. (c) Venn diagrams indicate the overlap between DEGs in (b) for control versus heat stressed samples. (d) Clustered gene expression heat map. (e) Gene ontology enrichment (fold change > 2, FDR < 0.01) of the up-regulated DEGs of SC7 vs. nSC7. (f) Normalized counts of reads for epigenetic regulators *H1*.*1, NRPD1A*, and *SWI3B*.

To understand which groups of genes form these five clusters, we performed a GO term analysis. Only clusters two and five revealed genes with significantly enriched annotations (Figure 3e). Interestingly, cluster 2 genes are only slightly induced by heat but are strongly repressed in non-stem cells and included genes for maintaining shoot development and the cell cycle. This suggests that stem cells maintain a cycling state and developmental programs during intense heat to facilitate recovery. Cluster 2 was also enriched for genes with negative regulatory roles. These genes comprised chromatin regulators, e.g., Histone *H1*.*1, NRPD1A* (a PolIV subunit), and the chromatin remodeling subunit *SWI3B* (Figure 3f); This indicates increased resilience of stem cells to chromatin decompaction, which occurs in nuclei of heat stressed plants (Pecinka et al., 2010; Dumur et al., 2019).

Genes of cluster 5 are characterized by strong down-regulation in stem cells and are enriched for stress response genes (Figure 3e), which again indicates that stem cells are maintaining their functions by suppressing stress-responsive pathways.

### 3.4. The influence of DNA methylation on heat response

Our subsequent analysis focused on heat-induced gene expression differences between the *wt* and the mutants with impaired asymmetric (*poliv*) or global (*ddm1*) DNA methylation. Comparing the immediate heat response in seedlings (Sd7) of *wt, poliv*, and *ddm1*, we identified more than 2000 up- and over 3000 down-regulated genes for all three genotypes (Wald test FDR < 0.05 and log2 fold change > |1|, Figure 4a). However, the number of DEGs in stem cell nuclei was lower, with only approximately 1100 up- and 2100 down-regulated genes, likely due to the absence of additive expression diversity originating from several cell types in seedlings.

**FIGURE 4.**
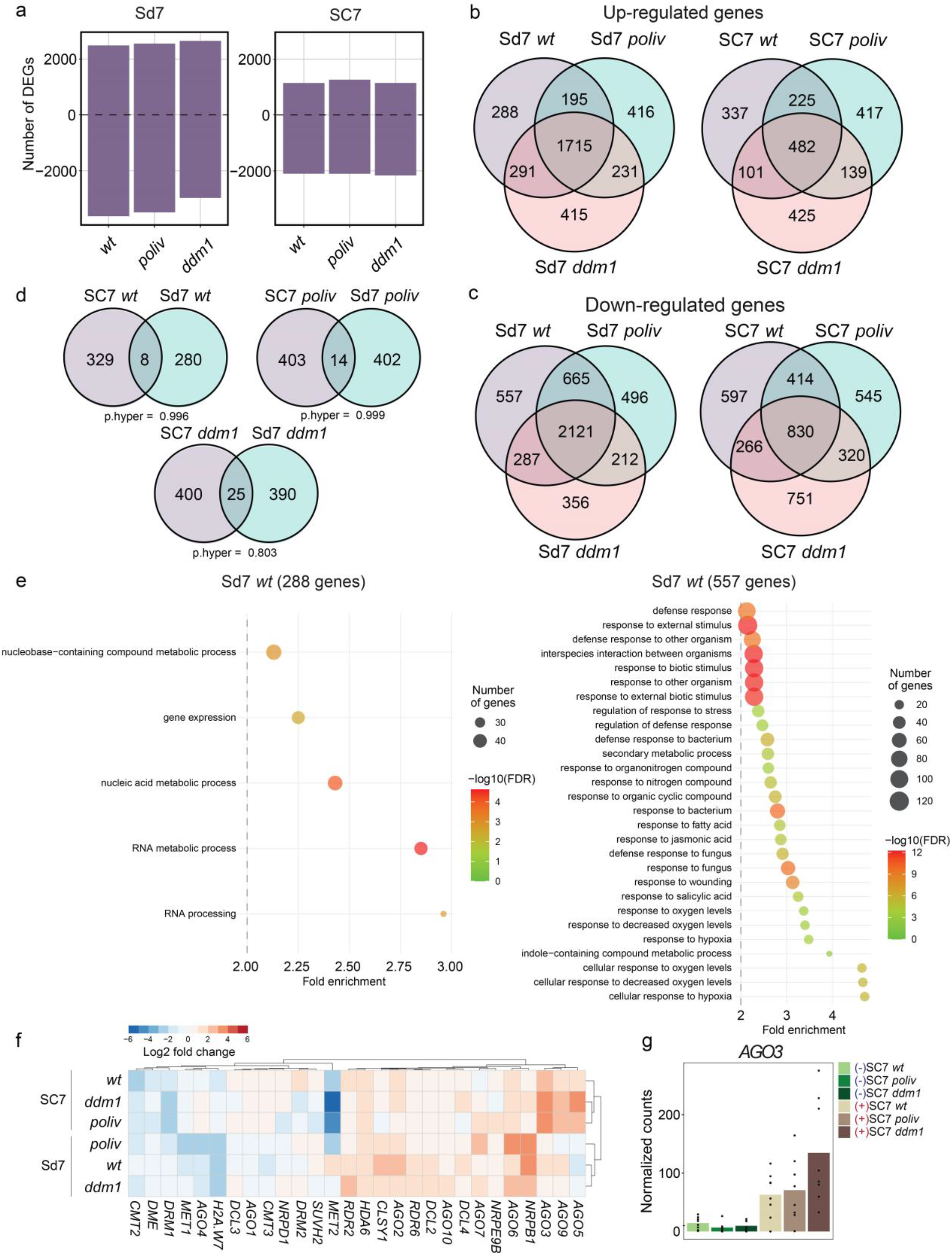
Heat responses in *poliv* and *ddm1*. (a) Number of differentially expressed genes (DEGs) under heat stress compared to control conditions. Three-way Venn diagram of up-regulated (c) and down-regulated (d) genes in *wt, poliv*, and *ddm1*. (e) Gene ontology analysis of distinctive up-regulated and down-regulated genes. (f) Expression heatmap of TE regulators in stem cells at D7 (SC7) and seedlings (Sd7). (g) Normalized counts of DCL4 in Sd7 under control (-) and heat stress (+).

Given the heat-hyposensitive phenotype of *poliv* and *ddm1*, we assumed that the expression of genes important for heat resilience and leaf necrosis prevention should change only in *wt* or in both *ddm1* and *poliv* seedlings. Therefore, we analyzed the overlap of gene expression changes (Figure 4b,c). The number of “private” DEGs for each genotype ranged between 288-425 (up-DEGs) and 496-751 (down-DEGs). These numbers were similar between stem cell nuclei and seedlings (Figure 4b), but their overlap was not significant (Figure 4d), underlining stem cell- and genotype-specific heat responses. To understand which functional group of genes could be causative for the difference in heat resistance, we performed GO-term analysis of *wt* private DEGs (288 up and 557 down), and DEGs present only in *poliv* and *ddm1* (231 up and 212 down). For *wt* private DEGs, we found 5 significant GO-terms for up- and 28 significant GO-terms for down-DEGs (Figure 4e). The GO-terms for the up-regulated genes described functions involved in RNA metabolism and contained among others *DCL4*, which is crucial for miRNA biogenesis and contributing to heat resistance (Figure 4f,g) (Popova et al., 2013). Interestingly, most genes of the down-regulated DEGs belong to stress response pathways, except for those of the heat stress response. This suggests that other defensive mechanisms were reduced to prioritize heat responses. GO-terms of DEGs changing in *poliv* and *ddm1* also included mostly stress-related categories (Supplementary Figure 2), and a loss of the homeotic balance between defense genes in these mutants could potentially contribute to increased heat sensitivity. Furthermore, enriched GO-terms of DEGs private to *ddm1* comprised categories of DNA damage and repair (Supplementary Figure 3b,e) and could indicate heat-induced genotoxic stress in *ddm1* caused by more frequent DNA breakage in decondensed chromatin or increased transposon activity.

As we expected to find up-regulation of TEs and observed upregulation of *AGO5* and *AGO9*-potential TE silencing factors, we also wanted to investigate which other host-counter defense genes become active. From an assembled list of genes involved in TE silencing, we identified *AGO3* (Figure 4g) as strongly induced by heat in stem cells in all three genotypes, but especially in *poliv* and *ddm1* (Figure 4f). AGO3 also binds to transposon-derived small RNAs of 24 nt lengths (Jullien et al., 2020; Zhang et al., 2016) and, together with AGO5 and AGO9, could confer protection against TE activity in stem cells under heat stress.

### 3.5 Persistent gene expression changes

At D28, three weeks after heat stress, gene expression differences were strongly reduced between mock- and heat-treated samples (Figure 5a). However, we still find 11, 8, and 39 genes up-regulated in stem cells (SC28) of *wt, poliv*, and *ddm1*, respectively (Figure 5a, Supplementary Table 1). The up-regulated genes of SC28 *wt* and *ddm1* contained *APA1* (*AT1G11910*), a gene that has been shown to confer drought tolerance when overexpressed (Sebastián et al., 2020). Most genes up-regulated in stem cells of *ddm1* were related to stress response (Table 1), including the heat shock transcription factor *HSFC1* (*AT3G24520*) (Supplementary Table 1), indicating that *ddm1* stem cells displayed an ectopic stress response three weeks after heat treatment.

**FIGURE 5.**
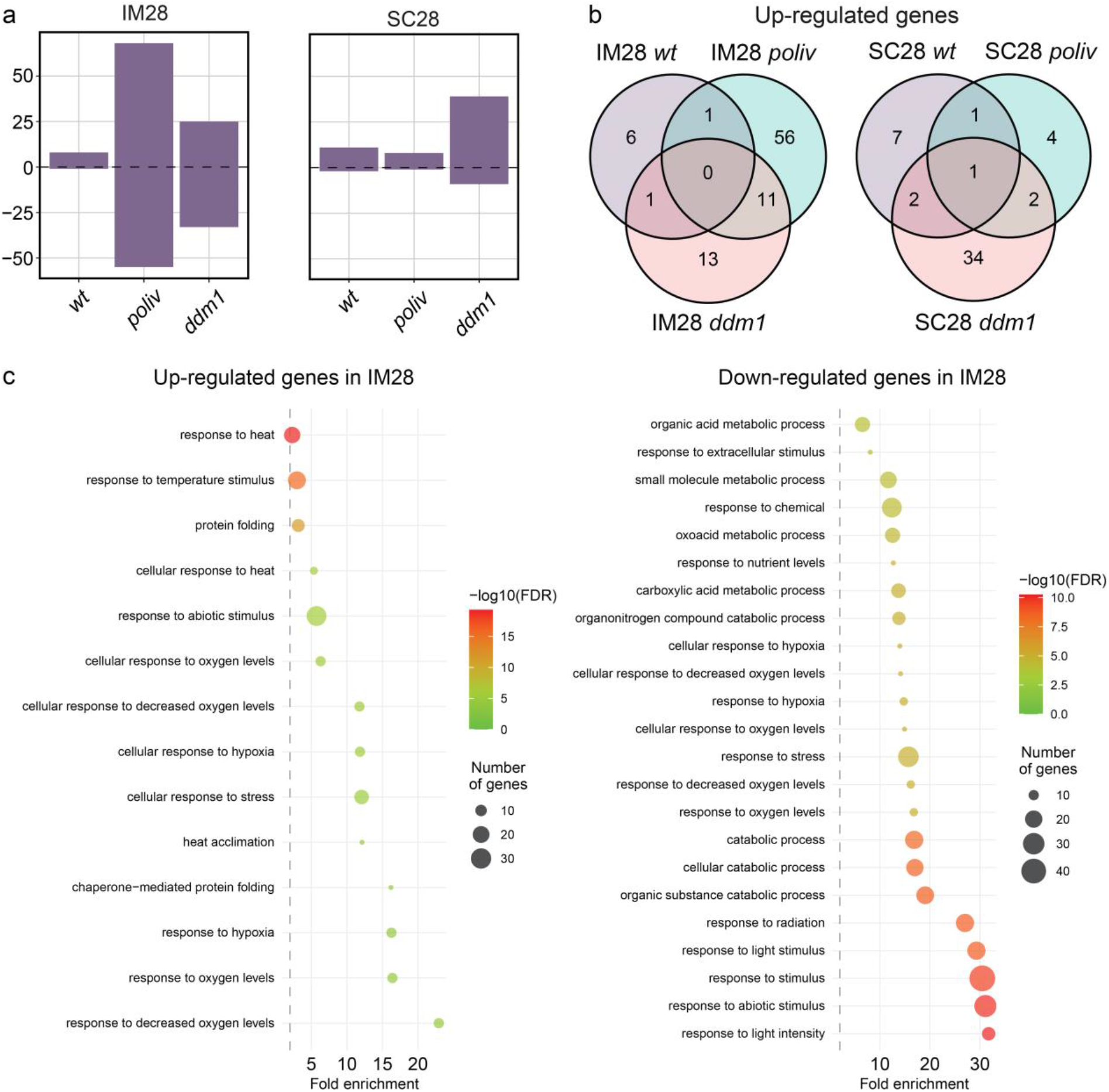
Persistent gene expression changes in *poliv* and *ddm1* after heat stress. (a) Number of differentially expressed genes (DEGs) in stem cell nuclei (SC28) and inflorescence meristem tissue (IM28) 21 days after heat stress. (b) Three-way Venn diagram of up-regulated genes in SC28 and IM in *wt, poliv*, and *ddm1*. (c) Gene ontology analysis of up-regulated and down-regulated genes in IM28.

**TABLE 1:**
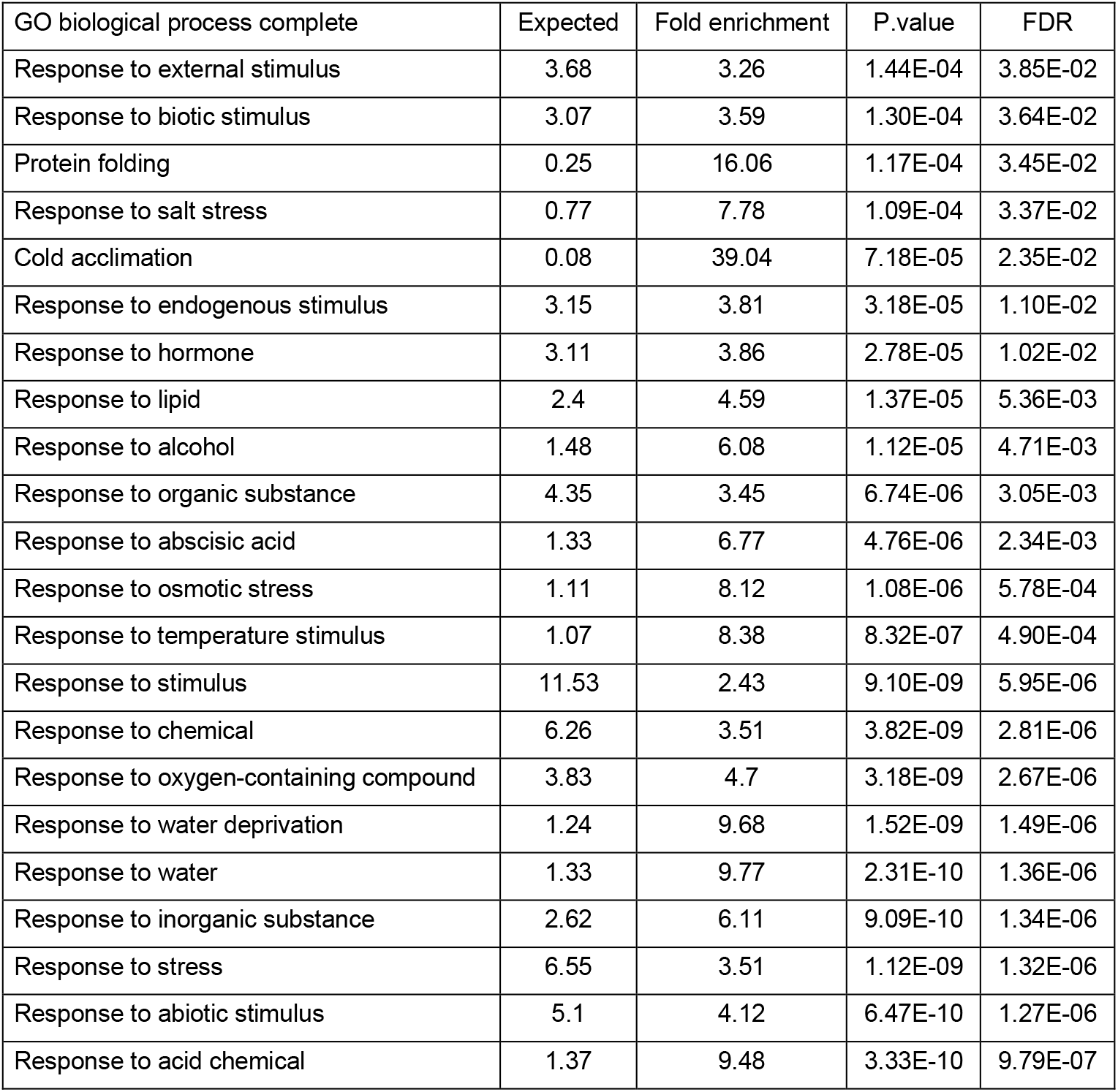
Gene ontology enrichment of up-regulated genes in SC7 *ddm1*.

Remarkably, we identified 25/68 up- and 33/55 down-regulated heat-induced DEGs in inflorescence meristems of recovered plants in *poliv* and *ddm1*, respectively (Figure 5a), in contrast to only 8 up- and 2 down-regulated genes in *wt*. Similarly to SC28, many of these up-DEGs are related to stress response pathways, including response to heat (Figure 5c). For *poliv*, 20 out of the 68 up-regulated genes are involved in heat stress response, including the heat shock transcription factor *HFSA2* (*AT2G26150*). HSFA2 has previously been shown to drive a transcriptional heat stress memory (Charng et al., 2007; Friedrich et al., 2021) and directly activates many heat stress responsive genes via a heat-response element (HREs) in the promoters of those genes (Schramm et al., 2006).

HSFA2 can also activate the transcription of TEs that contain the HRE element (Cavrak et al., 2014). This includes ONSEN and other TEs, and HSFA2 activity could be the main reason for the increase of TE expression upon heat stress (Pietzenuk et al., 2016).

### 3.6. Expression changes of transposons

Next, we focused our analysis on TE expression. First, we quantified the proportion of reads aligning to gene versus TE sequences (Figure 6a). In seedlings (Sd7), TE reads proportionally increased dramatically in all three genotypes (Figure 6a). We also observed increased TE transcripts in stem cell nuclei (SC7) of *wt* and *poliv* after heat treatment. In stem cells (SC) of *ddm1*, more than one quarter of all reads aligned to TEs, but surprisingly, this proportion decreased after heat treatment (SC7), and this decrease was still clearly visible after 21 days (SC28). This suggests that *ddm1* permits the expression of TE silencing factors that have a lasting effect on TE expression in stem cells under heat stress. On the contrary, we observed increased proportions of TE reads in heat-stressed *wt* and *poliv* samples 21 days after treatment (SC28 and IM; Figure 6a).

**FIGURE 6.**
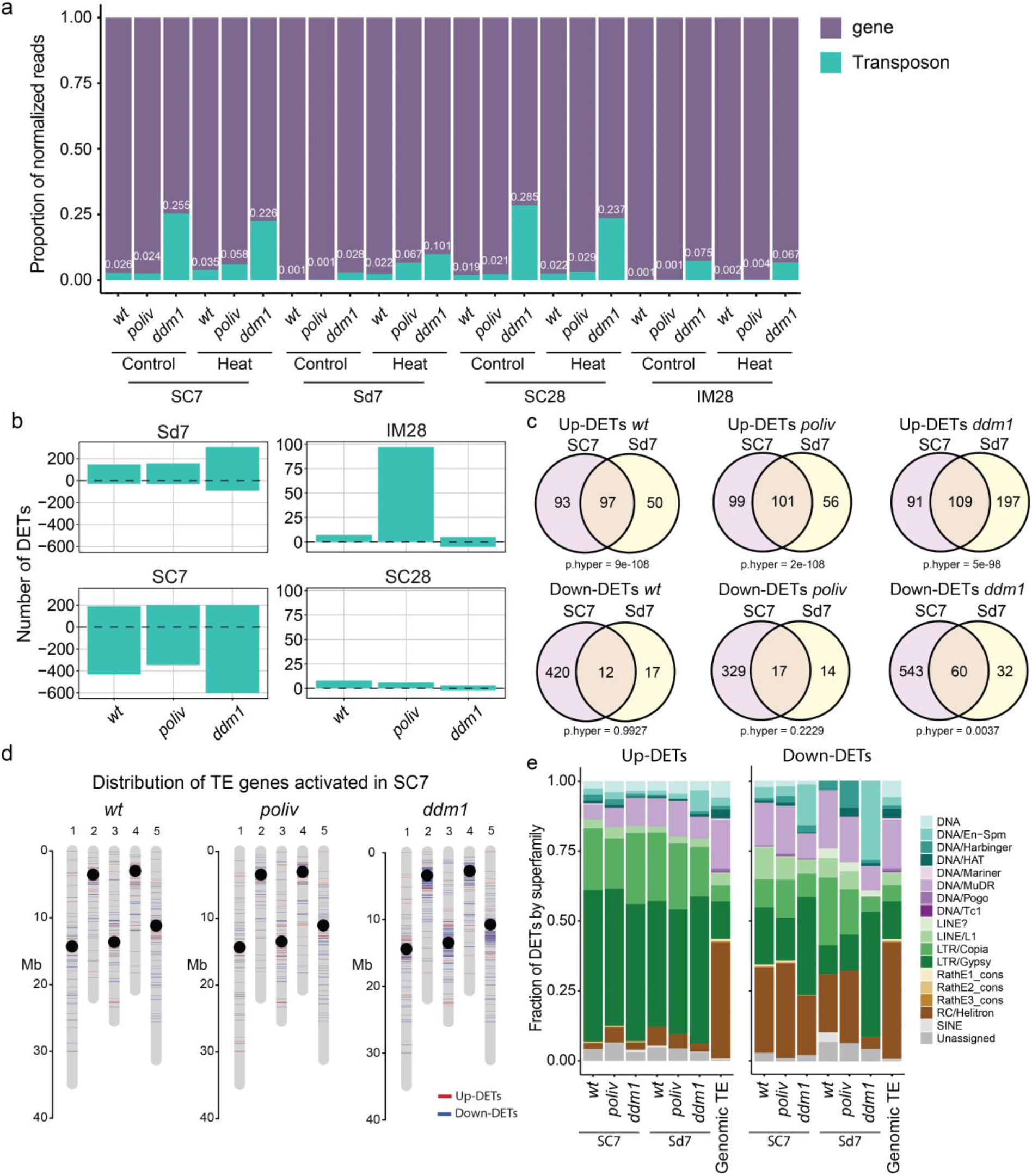
Expression of TEs in response to heat stress. (a) Fraction of genes and TEs by normalized counts. (b) Number of up- or down-regulated TEs by tissues: Sd: seedling, IM: inflorescence meristem, SC7: stem cell at D7, SC28: stem cell at D28. (c) Venn diagrams indicate overlaps of up- and down-DETs between SC7 and Sd. (d) Distribution of TE genes activated in SC7. (e) Fraction of DETs classified by superfamilies. Genomic TE: Proportion of the families in the reference genome.

To identify individual TEs that change expression upon heat, we calculated differentially expressed TEs (DETs Wald test FDR < 0.05 and log2 fold change > |1|) of heat-stressed vs. mock-treated samples for all genotypes (Figure 6b). The number of up-regulated DETs was similar between stem cell nuclei and seedlings, approximately 190 TEs (Figure 6b), and there was a significant overlap of heat-induced DETs in stem cells compared to seedlings (Figure 6c).

Also, 91 and 76 TEs were up-regulated independently of the genotype in seedlings and stem cells, respectively (Supplementary Figure 4a,b). Of those, 54 showed increased expression in stem cells and seedlings (Supplementary Figure 4c). These 54 TEs contained all 8 ONSEN copies and consisted mainly of LTR/Copia and LTR/Gypsy TEs (Supplementary Table 2). Twenty three of the 54 TEs were larger than 4 kb and could potentially be autonomous elements. Only stem cells showed a significant overlap for TEs with decreased expression after heat stress (Supplementary Figure 4b).

Up-regulated TEs in stem cells and seedlings were strongly enriched for LTR/Copia and LTR/Gypsy elements, but the superfamily distribution of down-regulated TEs was more similar to the genomic distribution and contained more DNA TEs (Figure 6e). Only TEs down-regulated in *ddm1* were also enriched for LTR/Gypsy elements (Figure 6e).

Loss of *DDM1* results in loss of DNA methylation, primarily at long heterochromatic TEs near centromeres (Zemach et al., 2013). Acute heat stress results in decondensation of chromocenters (Pecinka et al., 2010; Dumur et al., 2019), which consist of heterochromatic centromeres and pericentromeres. Therefore, we asked for the chromosomal localization of heat-induced up- and down-regulated DETs in our data set. Whereas up- and down-regulated DETs of heat-treated *wt* and *poliv* were found distributed over all parts of the chromosomes, DETs down-regulated in *ddm1* are enriched at the pericentromeres (Figure 6d), concordantly with the enrichment for LTR/Gypsy TEs (Underwood et al., 2017) (Figure 6e).

Intriguingly, we found 97 TEs still up-regulated in the inflorescence tissue of D28 *poliv* (Figure 6b), including two members of *ONSEN (ATCOPIA78*: *AT3TE92525* and *AT5TE15240*) (Figure 7a). We also detected highly increased expression (log2FC > 7.5) of four LTR/Gypsy, two DNA/MuDR, and two DNA TEs (Figure 7a). Most of these TEs belong to LTR/Gypsy and LTR/Copia elements (Figure 7b), and 65 out of the 97 TEs contained a potential HSFA2 binding motif (Supplementary Table 3). The presence of cytoplasmatic RNA in meristem samples and its absence in stem cell nuclei pools might explain the discrepancy in TE expression between the IM and SC28 in *poliv*. Extrachromosomal copies of retrotransposons (LTR/Gypsy and LTR/Copia) could also be templates for polymerases and could persist for long periods after the activating heat stress. The up-regulation of *HSFA2* in IM of *poliv* still after 3 weeks of heat exposure likely contributes to the activation of TEs.

**FIGURE 7.**
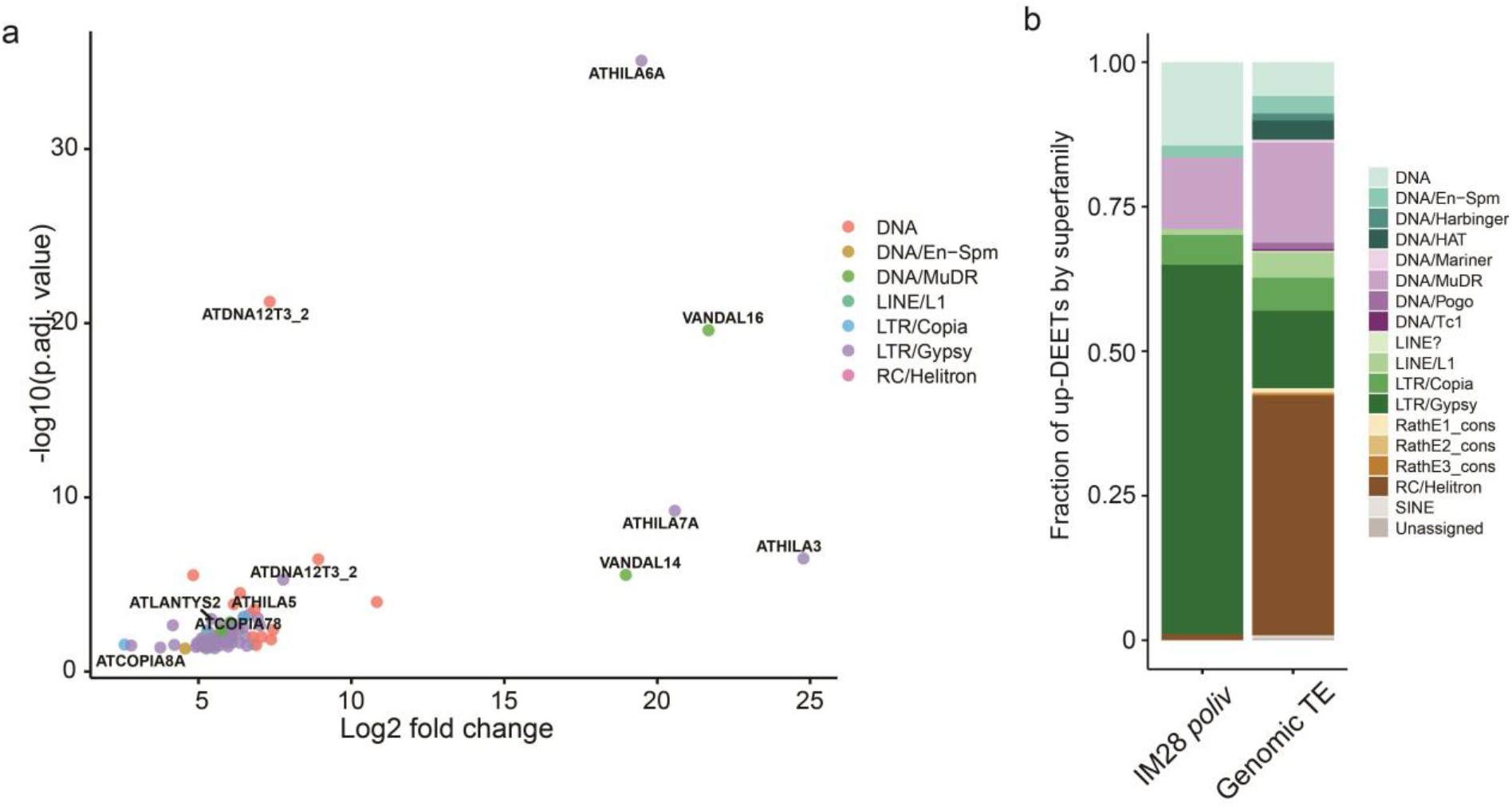
Persistant TE expression changes after heat stress. (a) Up-DETs of 97 TEs in IM28 *poliv*. (b) Fraction of TE superfamily of up-DETs in IM28 *poliv*.

### 3.7. Persistent DNA methylation changes in stem cells

DNA methylation changes at TE sequences can be associated with various stresses (e.g., Korotko et al., 2021; Wibowo et al., 2016). Furthermore, DNA methylation changes can last beyond several cell division cycles, be stably inherited, and influence phenotypes (Quadrana and Colot, 2016). A salt stress trigger can also induce maternally inherited DNA methylation changes (Wibowo et al., 2016). To be heritable to the next generation, DNA methylation changes should pass through the stem cells that form the germ line. Therefore, we analyzed DNA methylation of heat- and mock-stressed *wt, poliv*, and *ddm1* SAM stem cells three weeks after treatment. Large-scale DNA methylation patterns were similar between heat- and mock-treated samples (Figure 8a). As expected, TE DNA methylation in *ddm1* stem cells was low in all sequence contexts, and in *poliv*, CHG and CHH methylation was reduced (Figure 8b). However, we observed substantial differences in DNA methylation levels at TEs of heat-stressed stem cells, especially in the CHG context (Figure 8b). Heat induced a decrease of CHG TE methylation in *wt* but an increase in *poliv*. Interference of heat-induced chromatin decondensation with the activities of CMT2, CMT3, and RdDM could be the reason for this.

**FIGURE 8.**
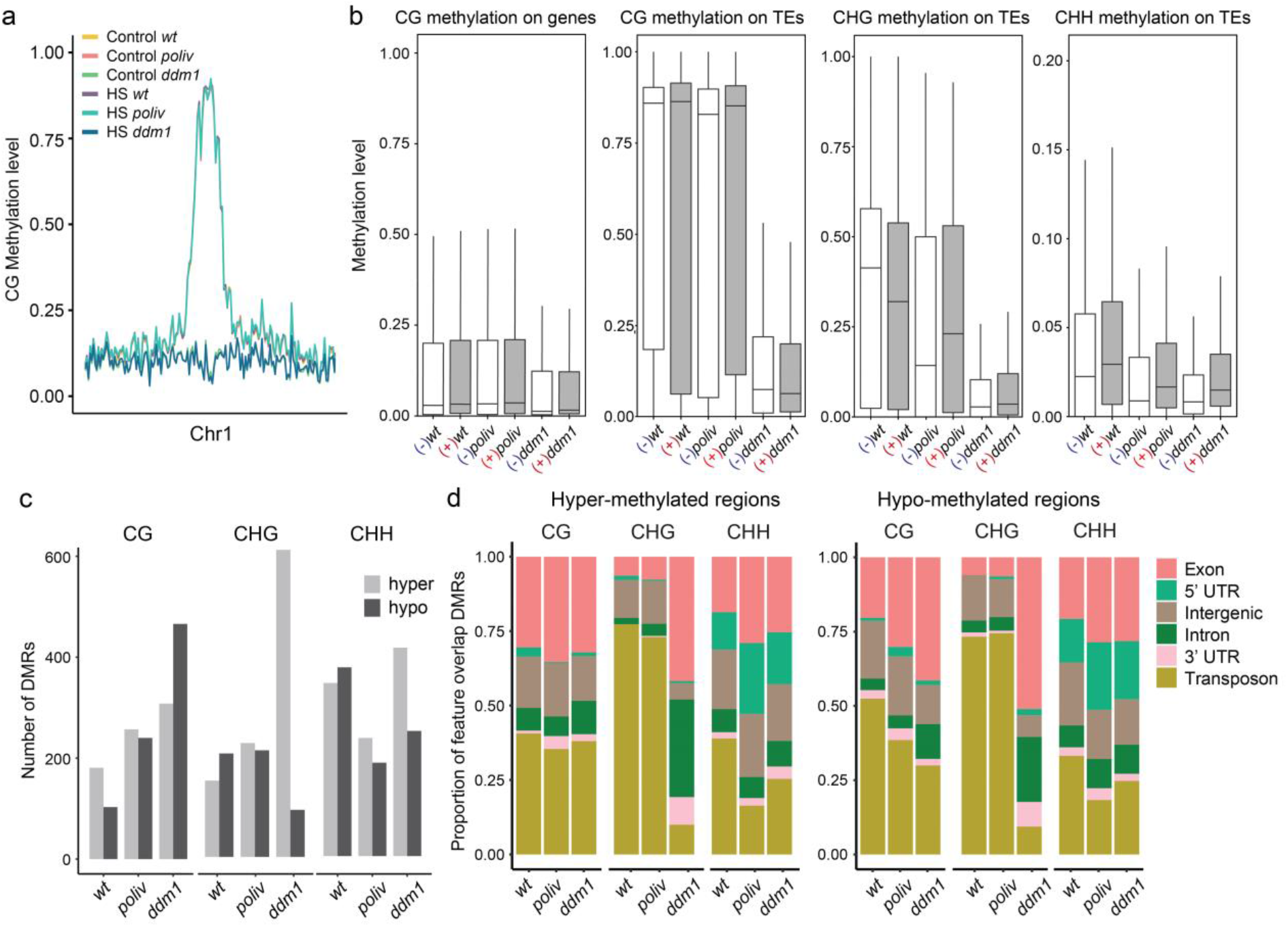
Heat stress triggers long-lasting DNA methylation changes in SAM stem cells. (a) CG methylation across chromosome 1, quantified in 200 kb windows. (b) Methylation levels on genes and TEs. (c) Differentially methylated regions at D28. (d) Hyper- and hypo-methylation sorted according to different genomic features.

Next, we identified differentially methylated regions (DMRs) employing a two-state Hidden Markov Model (HMM) (Hüther et al., 2022). This identified hundreds of heat-induced hypo- and hyper-DMRs in CG, CHG, and CHH contexts in all three genotypes (Figure 8c). *wt* showed more hyper-than hypo-DMRs at CGs but less in CHG and CHH contexts. *poliv* showed similar numbers of hyper- and hypo-DMRs in CG and CHG contexts but slightly fewer hypo-DMRs in the CHH context. Interestingly, *ddm1*, with generally low methylation levels in all sequence contexts, displayed the highest numbers of heat-induced DMRs, with less hyper-than hypo-DMRs in CG, but considerably more hyper-than hypo-DMRs in CHG and CHH contexts. Most DMRs overlapped with transposons or genic, a few with intergenic sequences (Figure 8d), indicating potential functional relevance. CG-DMRs overlapped mostly with transposon or exonic sequences. CHH-DMRs overlapped additionally to larger proportions with 5’UTR. Most DMRs in the CHG context overlapped with transposons in *wt* and *poliv*. This contrasts with *ddm1* CHG-DMRs, which overlap mainly with exonic and intronic sequences.

Although we could not find significant overlaps of DMRs with genes deregulated at D28 (Supplementary Table 4), we found 4 hypo-DMRs overlapping with 2 genes and 4 TEs with increased expression in IM28 *poliv*, and almost no hyper-DMRs overlapping with genes and TEs with reduced expression at D28.

Our analysis identified hundreds of DMRs that indicated lasting consequences of severe heat stress on DNA methylation in SAM stem cells for at least 3 weeks after stress. These DMRs can potentially be inherited or might be reset during germline differentiation. These DNA methylation changes seem to depend to some extent on the pre-existing DNA methylation state, as - in contrast to *wt* -, *poliv* showed an increase of CHG DNA methylation, and *ddm1* increased hyper-DMRs in CHG and CHH context after heat stress.

## 4. Discussion

Previous analysis of transcripts, DNA methylation, and transposon activity in the stem cells of the shoot apical meristem had provided evidence that here as part of the germline to the next generation, cell-specific and dynamic changes contribute to the epigenetic control of the genome (Gutzat et al., 2020; Bradamante et al., 2022). The present study addressed whether and how the underlying mechanism in stem cells are influenced by heat stress, previously shown to activate transposons. We analyzed stem cells of wild-type plants and two mutants impaired in two different pathways with an impact on DNA methylation (*ddm1* and *poliv*) for potential changes triggered by heat stress and for their maintenance beyond the stress exposure.

As expected, heat stress profoundly affected the transcriptome in all samples, mainly on stress-related genes and genes involved in nucleic acid metabolism, with differences between wt vs. *poliv* and *ddm1*. This could explain the hypersensitivity to severe heat stress common to both mutants. This contrasts previous results (Popova et al., 2013), where *ddm1* and *poliv* were not identified as sensitive to heat but several RdDM pathway mutants. However, heat stress in that study was applied on soil-grown mature plants and does not allow a comparison with the data from *in vitro*-grown young seedlings here. The transcriptome analysis needs to be refined and extended to investigate whether the heat sensitivity in DNA methylation mutants has a common basis. Still, a correlation of DNA methylation differences among geographic ecotypes with temperature (Dubin et al., 2015) might speak for this connection. However, the epigenetic control of heat tolerance in plants is complex, and much more is known about the role of other chromatin features (reviewed in Perrella et al., 2022).

Using a single-cell sequencing method for bulks of nuclei allowed us to study heat responses of shoot stem cells, which are crucial for post-embryonic development and hence essential for understanding developmental responses to heat stress. Our analysis indicates that after heat stress, cell cycle, and developmental programs are less suppressed in stem cells than somatic cells. This could allow a fast resumption of growth or organ regeneration after severe heat stress and agrees with an autonomous heat stress memory (Olas et al., 2021). We also found a strong heat-triggered up-regulation of *AGO3*, and further increased expression of *AGO5* and *AGO9* in stem cells of all three genotypes. All three AGOs are loaded with TE-derived sRNAs, but the respective mutants show little TE de-repression (Jullien et al., 2020; Bradamante et al., 2022). Either they can functionally complement each other, or their role for TE repression becomes only evident during heat- or other stresses, which are known to increase TE activity (Gutzat and Mittelsten Scheid, 2012; Dubin et al., 2018).

The retrotransposon ONSEN with its heat-responsive motif in the LTR (Cavrak et al., 2014) is a well-established indicator of lost epigenetic control under temperature stress, but despite strong accumulation of extrachromosomal DNA in heat-treated *wt* and *poliv*, its mobilization and integration into new genomic locations was only observed in *poliv* (Ito et al., 2010). This can be connected to our observation that increased levels of ONSEN and other heat-inducible TEs persisted in inflorescence meristems of *poliv* more than 20 days after heat exposure. Correspondingly, we also found increased and lasting expression of HSFA2, a heat response factor binding to the LTR of ONSEN and crucial for its heat-triggered induction. The prolonged “availability” of ONSEN in stem cells in *poliv*, but not in *wt* could be one reason for the lack of new insertions if the RdDM pathway is intact and the window for integration events is open only late in development. However, the role and mechanism of this persistent up-regulation have yet to be determined, as we could not find differences in DNA methylation in any of the up-regulated heat response factors or TEs. A recent study has shown that the 3D chromatin organization is also important for heat-induced transposon activation (Sun et al., 2020). Furthermore, heat stress induced large-scale chromatin organization changes, with an involvement of HSFA1a, another heat response transcription factor, and modifies numerous interactions of regulatory sequences in tomato (Huang et a., 2023). Future studies are necessary to comprehensively capture the changes at all levels and distinguish transient effects from those with long-lasting and transgenerational consequences. However, the hundreds of heat-induced DMRs persisting at least in our experimental time range can be seen as good reason to explore if, and when during development, stress-induced DNA methylation and other epigenetic changes occur. Furthermore, our work is a starting point to study to which degree such epimutations are reset or not, random or directed to distinct loci of the genome, whether they can be cumulative, and if they lead to permanent gene expression changes with phenotypic consequences. Investigating this stem cell- and other cell-specific stress responses will significantly deepen our knowledge about developmental plant adaptation during the plant’s lifetime and on a population level.

## Acknowledgments

We sincerely thank the following colleagues: Mattia Donà, Zsuzsanna Mérai and Philip Wolff for manuscript revisions. We are grateful for the excellent support by the GMI/IMP/IMBA Biooptics facilities, the Next Generation Sequencing and Plant Sciences units of the Vienna BioCenter Core Facilities (VBCF), and the EHS unit of the VBC that provided safe working conditions during the Covid pandemic. We gratefully acknowledge financial support from the Austrian Science Fund (FWF I489, I1477 to O.M.S and I3687 to R.G.), Vienna Science and Technology Fund (WWTF LS13-057) and Cost Action 16212 “INDEPTH” to O.M.S., and the Plant Fellows program (EU FP7) to R.G. We also acknowledge financial support for sequencing experiments via INTERREG RIAT-CZ to R.G.

## Author contributions

V.H.N. and R.G. conceived and designed the study. V.H.N. performed experiments, and V.H.N. and R.G. analyzed data. V.H.N., O.M.S., and R.G. wrote the manuscript. All authors read and approved the manuscript.

## Data availability

All RNA-Seq and Bisulfite-Seq data of this study are available in the Gene Expression Omnibus database at https://www.ncbi.nlm.nih.gov/geo/, reference number GSE223915.

