## Supplementary Figures for "Heat stress response and transposon control in plant shoot stem cells"

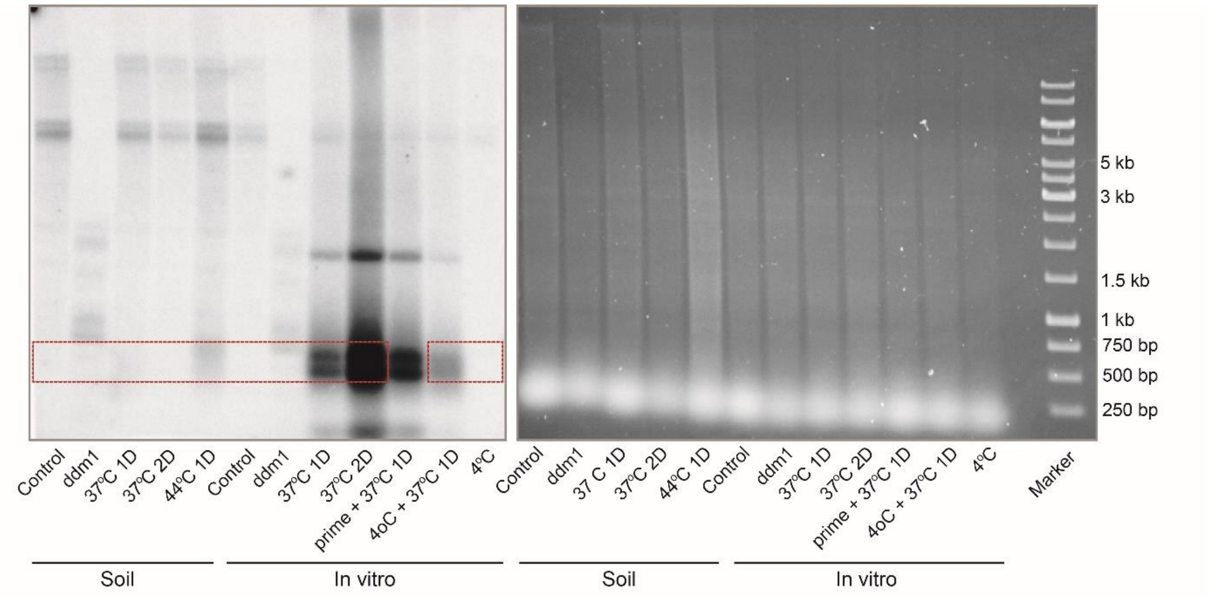


SUPPLEMENTARY FIGURE 1. Uncropped Southern blot. The red boxes indicate the sectors shown in Figure 1. The right side shows the loading of the corresponding gel before DNA transfer.

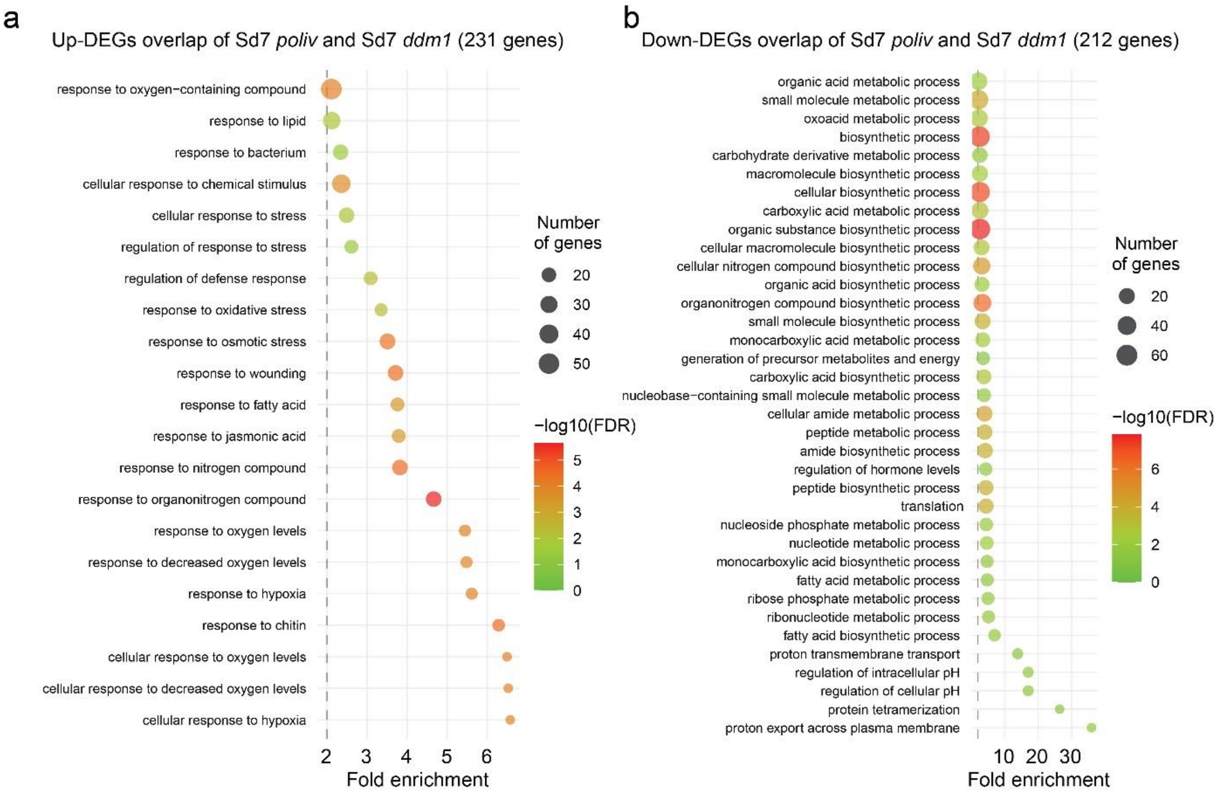


SUPPLEMENTARY FIGURE 2. Gene ontology analysis of overlaps between Sd7 *poliv* and Sd7 *ddm1* of upregulated (a) and downregulated (b) genes.

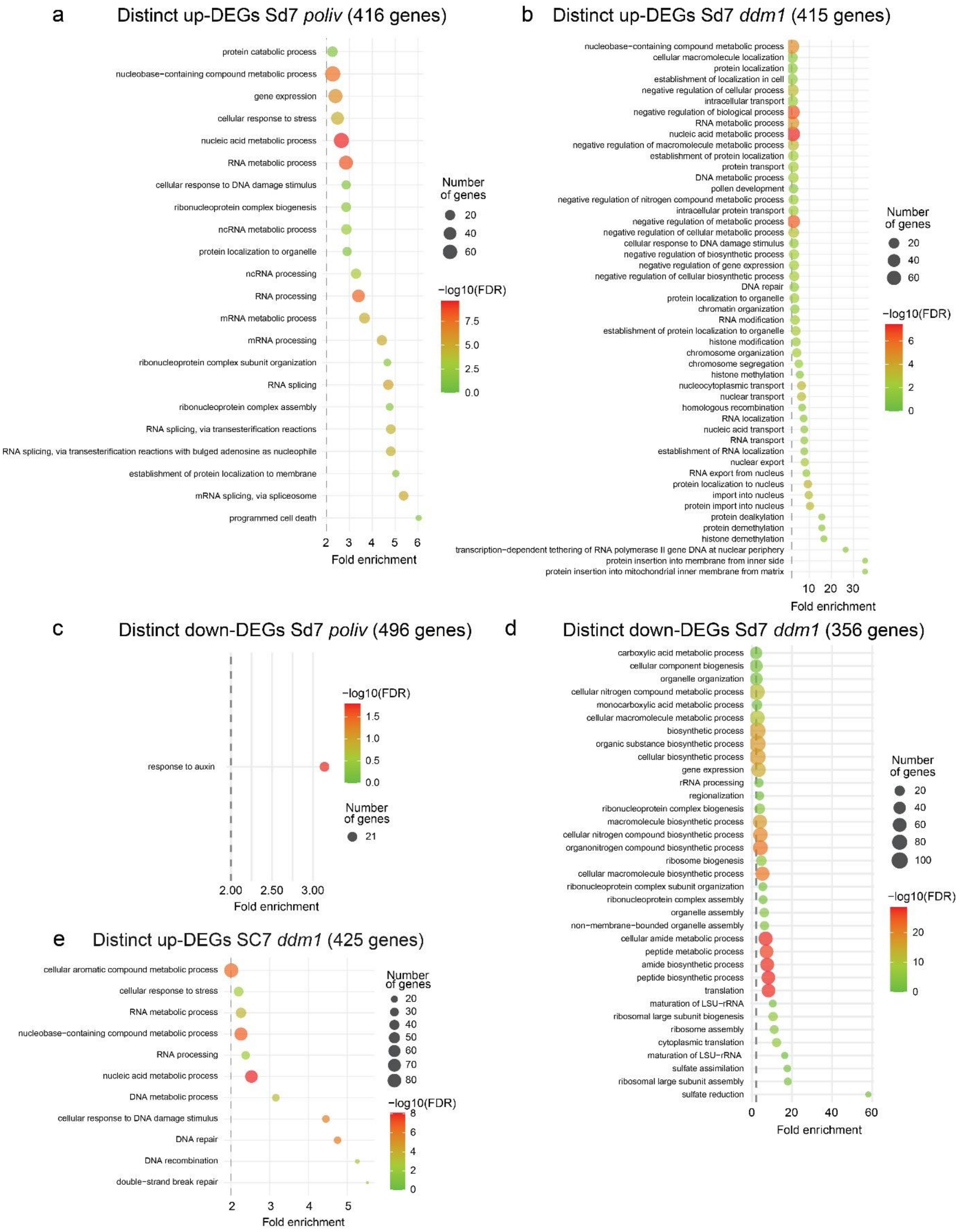


SUPPLEMENTARY FIGURE 3. Gene ontology analysis of distinctive upregulated genes of Sd7 *poliv* (a), Sd7 *ddm1* (b) and downregulated genes of Sd7 *poliv* (c) and Sd7 *ddm1* (d), and distinct upregulated genes of SC7 *ddm1* (e).

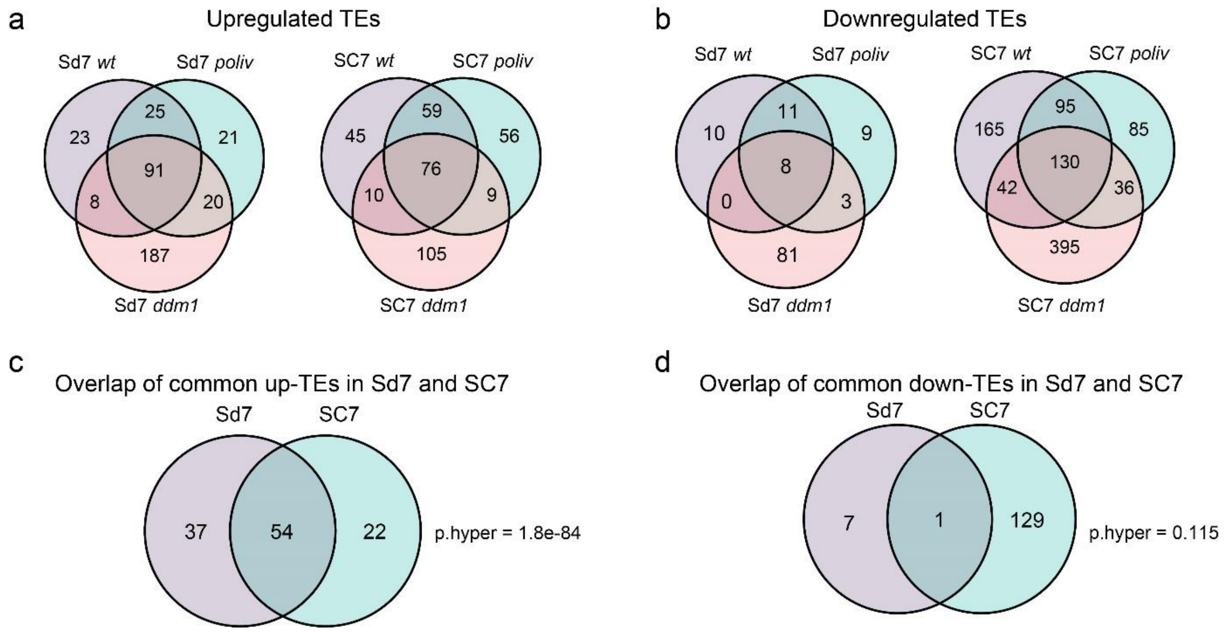


SUPPLEMENTARY FIGURE 4. Venn diagrams indicate common up- and down-regulated TEs. Three-way overlaps of upregulated (a) and downregulated TEs (b) in Sd7 and SC7. Overlaps of common sets in upregulated TEs (c) and downregulated TEs (d).
